# Evolution in spatiotemporal infection patterns of *Burkholderia* sensu lato lineages in the gut of *Riptortus pedestris*

**DOI:** 10.1101/2025.10.29.685444

**Authors:** Kota Ishigami, Antoine-Olivier Lirette, Yoshitomo Kikuchi

## Abstract

Many plants and animals form specific symbioses with microorganisms, relying on bidirectional interactions between hosts and bacteria. However, the knowledge about the evolution of symbiont traits enabling such specificity remains very limited. The bean bug *Riptortus pedestris* acquires *Caballeronia* from environmental soil and harbors it in its gut symbiotic organ. This bug-*Caballeronia* symbiosis is an ideal model to clarify the evolutionary process of symbiotic bacteria because members of their outgroups, such as *Paraburkholderia* and *Pandoraea*, can also colonize the host symbiotic organ but are outcompeted when co-inoculated with the native symbiont, *Caballeronia*. In this study, mechanisms underpinning the competitiveness of *Caballeronia* inside the insect gut were investigated. First, comparative microscopy revealed that *Caballeronia*’s success in the gut is largely attributed to its ability to migrate rapidly to the M4 region through chemotaxis, wherein a *cheA* insertion mutant showed significantly delayed infection speed and lower competitiveness against wild-type. In addition, *Paraburkholderia* and *Pandoraea* frequently formed biofilm-like aggregates in the midgut, which could delay their colonization. By contrast, *Caballeronia* formed no biofilm-like aggregates, at least inside the insect gut. This study reveals that *Caballeronia* symbiont has evolved traits like chemotaxis and reaction against AMP to establish an efficient and exclusive symbiotic relationship with their bean bug host. Although the genetic and molecular bases of the chemotaxis and the cell aggregates remain unclear, it is strongly suggested that the dynamic gain and loss of these traits enables *Caballeronia* to specifically associate with the insect host, *R. pedestris*.

**Importance:** *Riptortus pedestris*, a major soybean pest in East Asia, acquires symbiotic bacteria from the environment every generation, yet its gut is consistently and specifically colonized by *Caballeronia* species. The evolutionary traits that underlie this strong symbiotic specificity remain poorly understood. Here, we demonstrate that *Caballeronia insecticola* has acquired two key traits: positive chemotaxis toward the symbiotic organ and tolerance to host-derived antimicrobial peptides (AMPs). Using comparative colonization assays with wild-type, chemotaxis-deficient mutants, a non-symbiotic sister group, and their common ancestor, we show that these adaptations enable *Caballeronia* to outcompete other bacteria within the host gut. Our findings provide direct evidence that specific ecological and behavioral traits contribute to the establishment of exclusive symbiotic associations, shedding light on how symbiotic specificity can evolve even in horizontally transmitted systems. This work offers novel insights into the evolutionary dynamics of host-microbe interactions.

## Introduction

Most plants and animals form symbiotic relationships with microorganisms. Some symbioses establish specific and exclusive relationships. In such symbiosis, hosts have developed sophisticated partner choice mechanisms to acquire particular bacterial symbionts (1, 2). The symbioses achieve partner specificity through various host mechanisms (1–4), such as signal recognition and the secretion of antimicrobial peptides (5, 6). This specificity is not achieved solely by host traits; symbionts must also possess corresponding abilities to establish the relationship. In essence, specific symbiotic relationships are formed through bidirectional interactions rather than unidirectional processes. However, evolutionary insights into the abilities of symbiotic bacteria to establish specific symbiosis remain very limited. This limitation arises because systems with highly specialized symbiotic relationships often cannot associate with a wide variety of bacterial partners, including outgroup strains. As a result, it is experimentally difficult to investigate the stepwise processes by which such highly specific and stable partnerships have evolved.

The bean bug *Riptortus pedestris* provides an ideal model for studying the evolution of traits in symbionts for establishing specific symbiotic relationships. The stink bug is widely distributed in Southeast Asia and has a midgut divided into four sections, M1 to M4 (4). The M1 to M3 are involved in digestion, and infected with diverse microbial communities, while M4 is colonized by a stable, high-density population of a single microbial symbiotic species of the genus *Caballeronia*, (Fig. 1A) (7–9). In the M4 region, there are sac-like structures called crypts, densely packed with symbiotic bacteria. Although these symbionts are not essential for the survival of *R. pedestris*, they benefit the host insect by enhancing growth, reproduction, immune homeostasis, and pesticide resistance (5, 7, 8, 10–12). *Riptortus pedestris* acquires microbial partners from environmental soil each generation (8, 13). Despite the presence of diverse bacteria in the environmental soil, *R. pedestris* specifically retains *Caballeronia* bacteria in its symbiotic organ.

**FIGURE 1.**
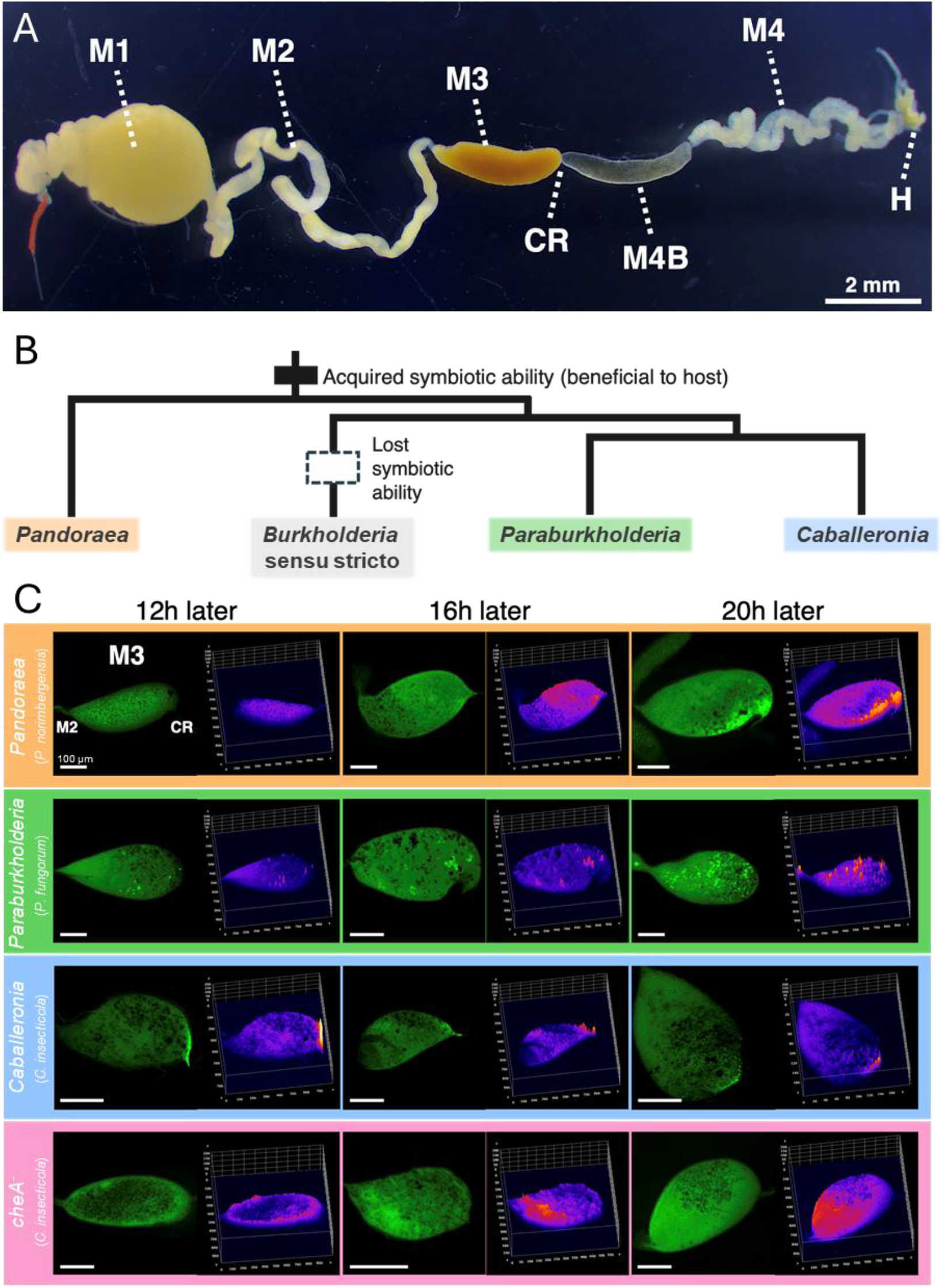
Bacterial distribution in the M3-Constricted region during infection. (A) Dissected midgut of *R. pedestris* infected with *Caballeronia insecticola*: M1, midgut first section; M2, midgut second section; M3, midgut third section; CR, constricted region; M4B, bulbous region prior to M4; M4, crypt-bearing midgut fourth section; H, hindgut. (B) Dendrogram of the phylogenetic relationships between Pandorea and genera of *Burkholderia* sensu lato. Note that all the genera except *Burkholderia* sensu stricto can colonize M4. The black box represents the evolution of the symbiotic ability; the white box represents the loss of symbiotic ability. (C) Visualization of the bacterial distribution in the M3 region during infection. Representative images 12h, 16h, and 20h after oral inoculation are shown. The bacteria are labeled by GFP. In each panel: left, a fluorescence microscopy image; right, a 3D surface plot of luminance. The luminance is interpreted as height for the plot.

Environmental bacteria ingested by *R. pedestris* are strictly selected by the “constriction region (CR)” located between M3 and M4, the lumen of which is only a few μm in diameter and is filled with mucus-like viscous substances. *Caballeronia* symbionts are capable of crossing the CR, whereas phylogenetically distant bacteria such as *Escherichia coli* cannot. Following infection by symbiotic bacteria, the CR region completely closes, preventing secondary infection (14, 15). Additionally, upon bacteria reaching the symbiotic organ, various antimicrobial peptides are specifically expressed, contributing to host specificity (5). Notably, while *R. pedestris* has these multiple mechanisms for partner choice, bacterial groups closely related to the genus *Caballeronia*, such as *Paraburkholderia* from the *Burkholderia* sensu lato group and *Pandoraea* from the out-group of *Burkholderia* sensu lato, can also pass through the sorting organ (CR), reach the symbiotic organ (M4), and proliferate and establish colonization in the symbiotic organ similarly to *Caballeronia* (Fig. 1B) (16). However, previous studies have shown that when *Pandoraea* or *Paraburkholderia* are orally ingested simultaneously with *Caballeronia*, the latter outcompetes the formers, ultimately dominating the symbiotic organ (16). What causes such a difference in the infectivity between the closely related bacterial groups remains unclear.

In this study, we closely observed the initial infection process of three bacterial genera: *Caballeronia, Paraburkholderia*, and *Pandoraea*, and elucidated quantitative differences in infectivity, such as the time taken to reach the M4 symbiotic organ and the invasion rate into the lumen of M4 crypts. Detailed observations revealed that *Caballeronia* reaches and passes through the CR faster than other bacteria, which enables *Caballeronia* to colonize and proliferate in the M4 symbiotic organ first, suggesting that chemotaxis plays a pivotal role to ensure the specific association. Furthermore, our observations demonstrated that *Pandoraea* and *Paraburkholderia* show delayed infection and colonization by forming biofilm-like aggregates within the midgut. These findings strongly suggest that the gain of chemotaxis and the loss of biofilm in the *Caballeronia* lineage are crucial for establishing specific symbiosis with the bean bug host, providing a novel insight into how symbiotic traits could evolve in symbiont lineages.

## Materials and Methods

### Insects and Bacterial Strains

The *Riptortus pedestris* strain used in this study was originally collected from a soybean (*Glycine max*) field in Tsukuba, Ibaraki Prefecture, Japan, and has been maintained in the laboratory for over ten years. The insects were reared in Petri dishes (90 mm diameter, 20 mm height) at 25°C under long-day conditions (16 hours light, 8 hours dark), and were provided with soybean seeds and distilled water containing 0.05% ascorbic acid (DWA). For the experiments, GFP-labeled strains of *Caballeronia insecticola, Paraburkholderia fungorum*, and *Pandoraea norimbergensis* were used. In addition, a chemotaxis-deficient strain (*cheA* insertion mutant) was employed. The *cheA* mutant strain used in this study was originally generated by Ohbayashi et al. (2015) as a transposon insertion mutant constructed by using the Mini-Tn5 system (17). The *cheA* gene encodes a histidine kinase that plays an essential role in bacterial chemotaxis (18). All symbiotic bacteria used in this study, including the *cheA*^*-*^, are listed in Supplementary Table S1. The *cheA* insertion mutants was labeled with GFP using the Tn7 mini-transposon system as previously described (7).

### Oral Administration of Bacteria

Each symbiotic bacterial strain was cultured at 30°C in YG medium (with 30 μg/ml kanamycin for GFP-labeled strains) on a rotary shaker (150 rpm) until the early logarithmic growth phase. The bacteria were harvested by centrifugation, resuspended in DWA, and adjusted to a concentration of 10^4 cells/μl. After the first instar nymphs molted into the second instar, DWA was removed from the rearing containers to prevent the nymphs from drinking water overnight. Then, 1 μl of the adjusted bacterial suspension was provided to each second instar nymph, and individuals that consumed the suspension within one hour were used in subsequent experiments. After infection, the insects were supplied with DWA and reared.

### Dissection and Observation of Infection Dynamics

Insects that orally ingested bacteria were dissected in phosphate-buffered saline (PBS: 137 mM NaCl, 8.1 mM Na2HPO4, 2.7 mM KCl, 1.5 mM KH2PO4 [pH 7.4]) using tweezers under a dissecting microscope (S8APO, Leica, MZ FZ III, Leica). The midgut was then observed on a glass slide with PBS using a stereomicroscope (M205 FA, Leica, Germany) and photographs of the dissected tissues were taken with a digital camera (EC3, Leica). To examine the speed at which symbiotic bacteria reach the symbiotic organ, insects were dissected at regular intervals (0h, 4h, 8h, 12h, 16h, 20h, 24h, 40h) after oral inoculation, and GFP signals in the symbiotic organ (M4B, M4) was observed. Presence of bacterial in the symbiotic organ was considered positive if even a single GFP expressing bacterial cell was observed. The acquired data were statistically analyzed using the GraphPad Prism 10.1.0 (GraphPad Software). To compare the infection dynamics among bacterial strains, we examined differences in infection rates at each time point using Fisher’s exact tests for all pairwise combinations of strains, followed by Bonferroni correction for multiple comparisons. This analysis allowed us to evaluate how rapidly each bacterial strain reached the symbiotic organ. Additionally, to investigate bacterial localization in M3 before reaching the symbiotic organ, M3 was observed at 12h, 16h, and 20h post-ingestion.

Furthermore, to clarify which crypt in the M4 symbiotic organ the infection starts in, GFP signals in the crypt lumen were measured at each time point (12h, 16h, 20h, 24h, 32h, 40h, 120h). Five individuals per time point for each bacterial strain were dissected and counted. Crypts with observed GFP signals were considered positive. The total number of crypts at both ends of each symbiotic organ was counted, and numbered 1, 2, 3, etc., from the M3 side (*i*.*e*. from anterior to posterior). The value of the crypts number/total crypts number was used as the relative position within the symbiotic organ, where values close to 0 indicated the front part (M4B side) and values close to 1 indicated the hindgut side of the crypts.

### Competitive Infection Assay in the Gut Symbiotic Organ

Logarithmic phase cells of the wild-type strain and a *cheA*^*-*^ of *C. insecticola*, along with competitor bacteria (*Paraburkholderia fungorum* or *Pandoraea norimbergensis*), were suspended in distilled water, mixed, and adjusted to a concentration of 5,000 CFU/μL. For each assay, newly molted second instar nymphs of *R. pedestris* were individually fed 1 μL of the bacterial suspension as described above. The insects were reared at 25°C under long-day conditions with soybean seeds and distilled water. Five days after inoculation, symbiotic organs were dissected from each insect and homogenized in sterile distilled water. The suspensions were serially diluted and plated on YG agar medium containing antibiotics. After 3 days of incubation at 25°C, the colonies of SBE and competitor bacteria were counted. Competitive Index (CI) values were calculated using the formula [(output *C. insecticola* or *cheA*^*-*^ CFUs/input *C. insecticola* or *cheA*^*-*^ CFUs) / (output competitor CFUs/input competitor CFUs)] and statistically evaluated by the 1-sample t test (against CI = 1.0). Wild-type and chemotaxis mutant strains of *C. insecticola* were selected with rifampicin, and *Paraburkholderia fungorum* and *Pandoraea norimbergensis* were selected with streptomycin. The acquired data were statistically analyzed using the GraphPad Prism 10.1.0 (GraphPad Software).

### Effect of polymyxin B on bacterial growth

To evaluate the susceptibility of each bacterial strain to polymyxin B, an antibiotic that attacks bacterial cell membranes, a growth inhibition assay was performed using a 96-well microplate format. The three bacterial strains were pre-cultured overnight in YG medium, and then diluted with fresh medium to OD600nm = 0.3 and sub-cultured until they reached OD600nm ≈ 1. The cells were pelleted by centrifugation, resuspended in fresh medium, and diluted to OD600nm = 0.05. The cell suspensions were distributed into 96-well plates, one column per test strain, with 100 μL per well, except for the first column. The first column received a mixture of YG cell suspension mixed with Polymixin B dissolved in water, so that the final volume in each well would be 200 μL with a concentration of Polymixin B of 50, 25, or 12.5μg/mL depending on the column. Two-fold serial dilutions in subsequent columns were achieved by transferring 100 μL from one column to the next and mixing by pipetting up and down. The last column of the 96-well plate did not contain the peptide. This setup allowed testing of three polymyxin B (PMB) concentrations forming a concentration series of 50 μM, 25 μM, 12.5 μM, and control samples without PMB. The 96-well plate was incubated in a plate incubator (Synergy H1, Biotek). Growth of cultures in the wells was monitored by measuring OD600nm, with data points collected every 0.5 hours over 21 hours. The plates were incubated at 28°C with double orbital shaking at 200 rpm. The assay was performed four times.

### Measurement of *in vitro* Biofilm Formation

To quantify bacterial biofilm formation, crystal violet staining was performed. Freshly grown cell cultures were diluted in YG medium to OD600 = 0.3. Next, 2 μl of the cell suspension was inoculated into 198 μl of YG medium in 1.5 mL polypropylene tubes and incubated at 25°C 5days. Biofilm formation activity was assessed using a slightly modified crystal violet staining method (19). The liquid culture was carefully removed from the tubes, which were then rinsed once with distilled water. The biofilm formed on the inner wall of the tubes was stained with 0.1% crystal violet solution at room temperature for 20 minutes. After removing the free dye by washing twice with distilled water, the dye attached to the biofilm was extracted with 200 μl of 99% ethanol and quantified by measuring absorbance at 600 nm.

## Results

### Time to Reach the Symbiotic Organ and Chemotaxis

Observation of bacterial distribution in the M3 region after inoculation revealed that *C. insecticola* was concentrated near the CR twelve hours after oral intake (Fig. 1C). In contrast, this distribution bias was not observed at any time point for the chemotaxis mutant strain (Fig. 1C). Similarly, *Paraburkholderia fungorum* did not show such distribution bias towards the CR, and the bacteria were found clustered within the M3 region. *Pandoraea norimbergensis* showed a distribution bias near the CR, but this was observed 20 hours after ingestion, which was slower compared to *C. insecticola*.

Second instar nymphs of *R. pedestris* were fed with each bacterial strain, and their symbiotic organs were observed under a microscope to monitor bacterial colonization of the symbiotic organ (Fig. 2A and B; Fig. S1). Four hours after ingestion, both the wild-type *C. insecticola* and the chemotaxis mutant strain, *cheA*^*-*^, had already begun to colonize the M4 region, whereas *Paraburkholderia fungorum* and *Pandoraea norimbergensis* showed no detectable colonization at this stage (Fig. 2A; Fig. S1). The colonization rate of *C. insecticola* wild type and the *cheA*^-^ at 4 h was significantly higher than that of *Paraburkholderia fungorum* and *Pandoraea norimbergensis* (*P* < 0.05, Fisher’s exact test with Bonferroni correction). At 8 h post-ingestion, *C. insecticola* had colonized the M4 region in almost all individuals (95%, 19/20), which was significantly higher than the colonization rates of all other strains (*P* < 0.05, Fisher’s exact test with Bonferroni correction). The colonization pattern of the *cheA*^*-*^ mutant was comparable to that of *Pandoraea norimbergensis*, whereas *Paraburkholderia fungorum* showed the slowest progression among all tested strains (Fig. 2A; Fig. S1). By 16 h post-ingestion, the colonization rates of all strains except *Paraburkholderia fungorum* were significantly higher than that of *Paraburkholderia fungorum* (*P* < 0.05, Fisher’s exact test with Bonferroni correction). Finally, by 40 h post-ingestion, *Paraburkholderia fungorum* had also reached a 100% colonization rate in the M4 region, indicating that it eventually colonizes the organ but with a substantial delay compared to *Caballeronia* and *Pandoraea* (Fig. 2A; Fig. S1). These observations demonstrate that *C. insecticola* exhibits strong chemotaxis toward the M4 symbiotic organ, enabling rapid migration and colonization, while the *cheA*^*-*^ mutation results in delayed infection similar to that of the non-symbiotic *Pandoraea*.

**FIGURE 2.**
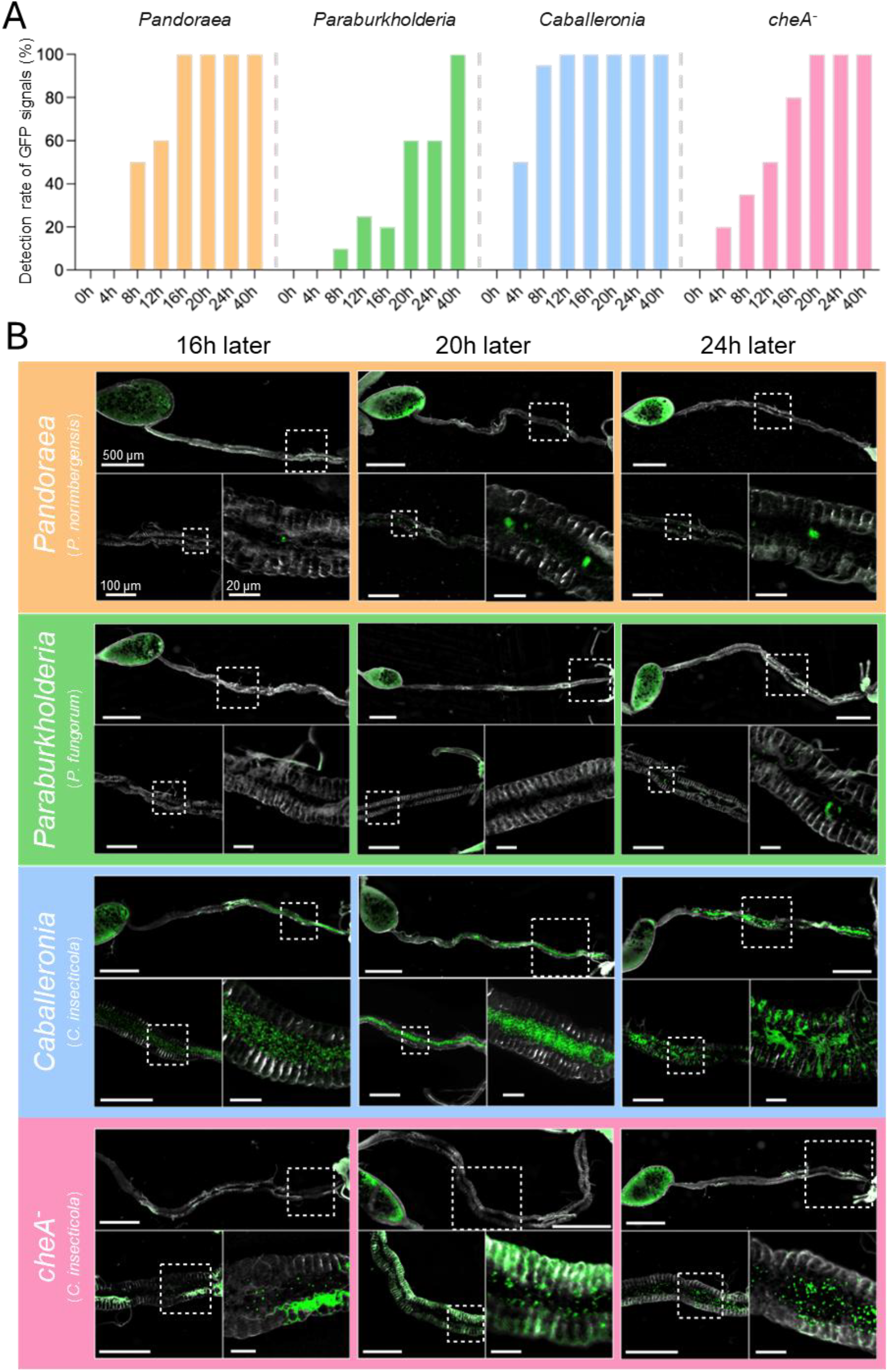
Time to reach the symbiotic organ. (A) The Time after inoculation at which GFP signals were detected in symbiotic organs. For each time point and each bacterium, twenty insects were investigated. If even a single bacterial GFP-signal was detected in the main duct of the M4 region, it was counted as “positive”. The Y-axis represents the proportion of positive insects. (B) Images of the GFP-labeled bacteria in M4 region.

### Infection Patterns after Reaching the Symbiotic Organ

Microscopic observations of the symbiotic organs post-ingestion revealed that bacterial proliferation occurred within the main duct before entry into the crypt lumen (Fig. 2B). This general pattern was visually consistent among all bacterial strains examined. To qualitatively describe infection dynamics within the symbiotic organ, we observed the timing and position of crypt colonization by each bacterial strain. *C. insecticola* was first observed invading the crypts approximately 16 hours after ingestion (Fig. 3). The initial infections were mainly seen in the posterior crypts, and this positional tendency was reproducibly observed at later time points. In contrast, the chemotaxis mutant began crypt invasion around 20 hours post-ingestion, but no clear spatial pattern of crypt colonization was apparent (Fig. 3). *Paraburkholderia fungorum* and *Pandoraea norimbergensis* showed delayed invasion, with bacteria appearing in the crypts after about 24 hours (Fig. 3). Interestingly, *Pandoraea. norimbergensis* tended to appear first in the anterior crypts, whereas *Paraburkholderia fungorum* was more often found in the posterior region, though this trend was less pronounced than in *C. insecticola*. These observations collectively suggest that chemotaxis affects both the timing and spatial progression of bacterial colonization within the symbiotic organ.

**FIGURE 3.**
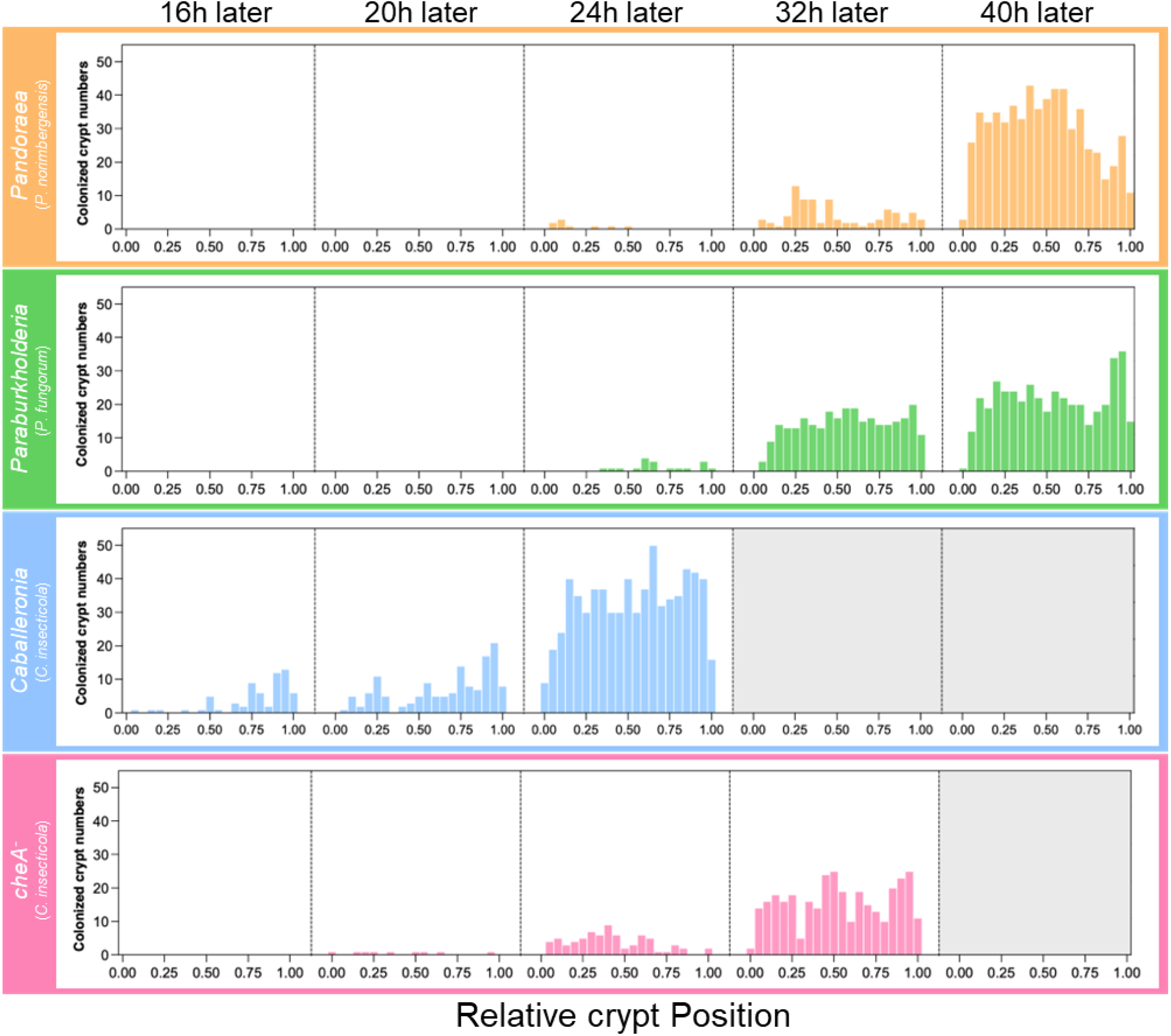
Colonization geography in crypts after reaching the M4 region. The Histogram mean numbers of crypts that are detected GFP signal of bacteria. X-axis indicates the relative crypt position from the M4B side (value 0) to the hindgut side (value 1.00). If even a single bacterial GFP-signal was detected in the crypts, it was counted as “colonized”. At each point and in each strain, five insects were investigated.

Observation of the symbiotic organs five days post-ingestion revealed that both *C. insecticola* and its *cheA*^*-*^ mutant strain had colonized most crypts within the M4 symbiotic organ (Fig. S2A and B). The colonization frequency of crypts by these strains was significantly higher compared to *Pandoraea norimbergensis* (*P* < 0.05, Mann-Whitney U test with Bonferroni correction). The colonization rate of *Paraburkholderia fungorum* was intermediate between *C. insecticola* and *Pandoraea norimbergensis* (Fig. S2A and B), with no significant differences with either. These results suggesting that chemotaxis is not crucial for the final colonization of crypts and indicate that *Pandoraea norimbergensis* is less efficient in crypt colonization.

### Importance of Chemotaxis in *in vivo* Competition

When wild-type *C. insecticola* was administered to *R. pedestris* along with *Pandoraea norimbergensis* or *Paraburkholderia fungorum*, the competitive index within the symbiotic organ after five days showed that *C. insecticola* was dominant (*P* < 0.05, 1-sample t test) (Fig. S3). However, when a *cheA*^*-*^ mutant of *C. insecticola* was co-infected with *Pandoraea norimbergensis* or *Paraburkholderia fungorum*, the chemotaxis mutant was unable to completely outcompete the other bacteria (Fig. 4). Furthermore, when both wild-type *C. insecticola* and the *cheA*^*-*^ mutant were administered to *R. pedestris*, the competitive index within the symbiotic organ indicated that the wild type dominated (*P* < 0.05, 1-sample t test) (Fig. 4). These results demonstrate that chemotaxis is crucial for competitive success within the symbiotic organ.

**FIGURE 4.**
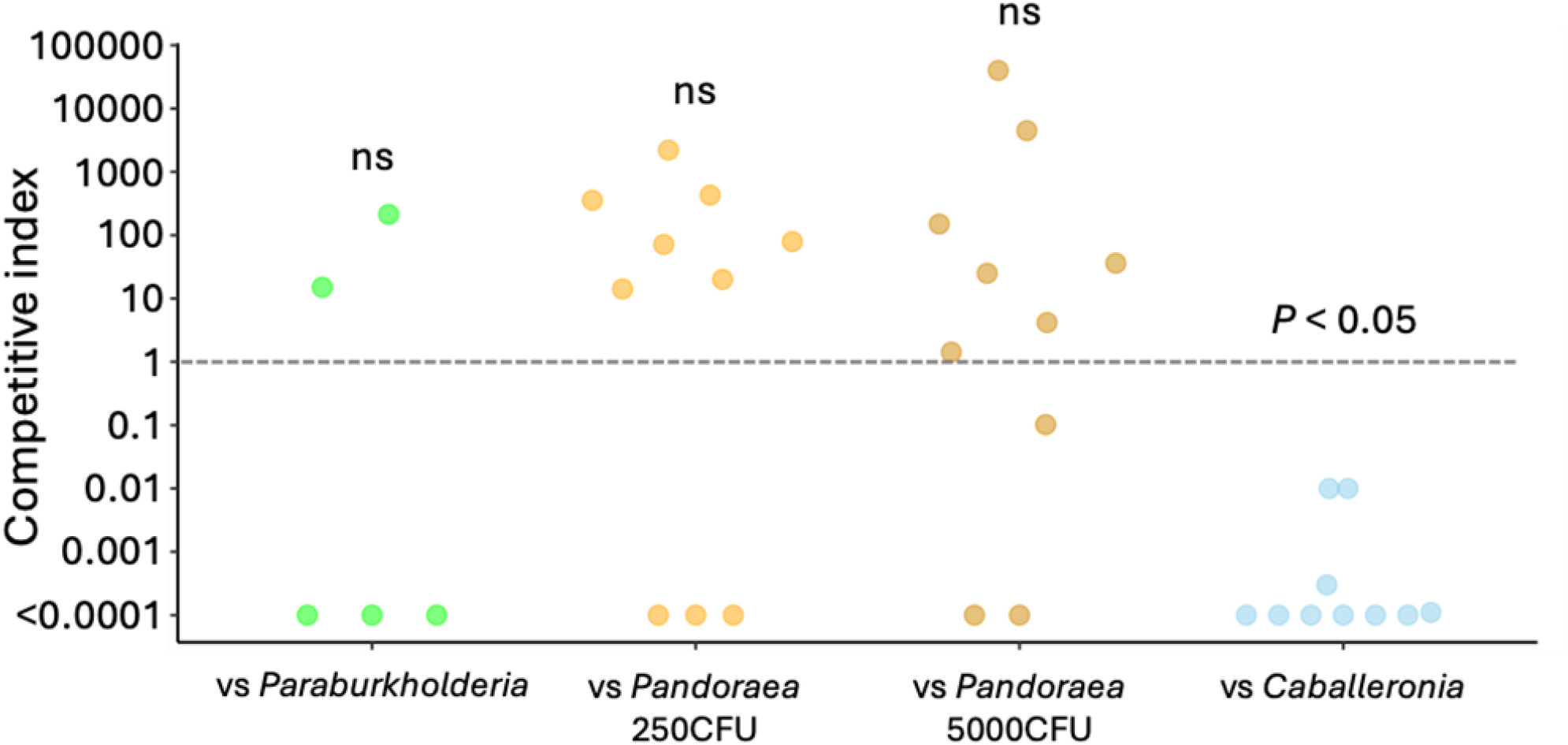
Competitiveness of chemotaxis mutant. Competitiveness of *cheA*^-^ within symbiotic organs. Competitive index values of were obtained by (output *cheA*^-^ CFU/input *cheA*^-^ CFU)/(output competitor CFU/input competitor CFU) and statistically evaluated by the 1-sample t test (against CI = 1.0).

### Biofilm Formation

Microscopic observations revealed that *Paraburkholderia fungorum* formed visible aggregates within the M3 region (Fig. 1C), whereas in M4, cells were mostly found either singly or in short chains (Fig. 5A). In contrast, *Pandoraea norimbergensis* did not show obvious aggregation in M3 (Fig. 1C) but exhibited distinct clusters within the main duct of M4 (Fig. 2 and 5A). *C. insecticola* cells were not observed forming such aggregates and appeared more dispersed throughout both M3 and M4. *Caballeronia* symbionts were observed moving and invading the crypt lumen more actively compared with *Pandoraea* and *Paraburkholderia* species (Supplementary movies S1-3). These observations suggest that forming aggregates (biofilm formation) is a crucial factor allowing faster and more efficient infection in the symbiotic organ.

**FIGURE 5.**
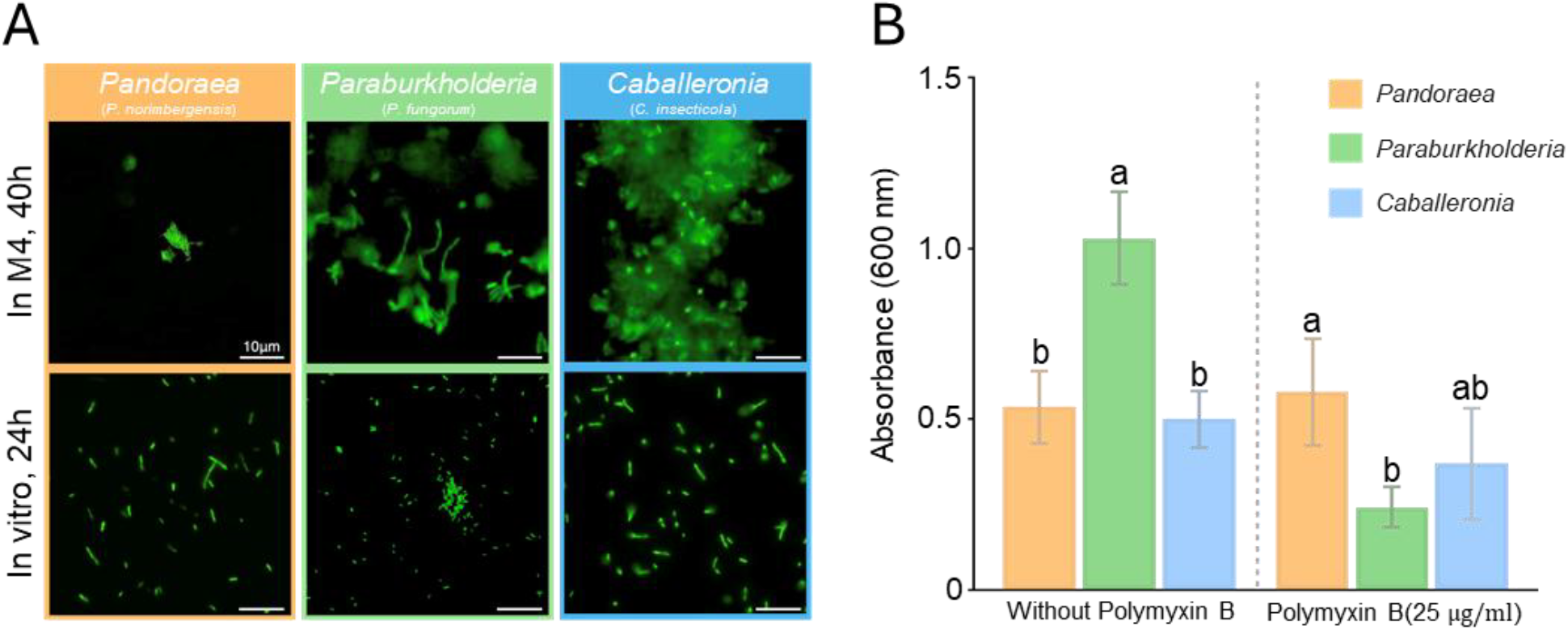
Properties of biofilm formation in *Caballeronia* symbiont and allied species. (A) *In M4* and *in vitro Pandoraea, Paraburkholderia* and *Caballeronia*. (B) Biofilm formation when Polymyxin B was added to in vitro culture. Biofilm formation was quantified by Crystal Violet staining. The different letters indicate statistically significant differences (P<0.05). The statistical significances were analyzed by The Mann-Whitney U test with Bonferroni correction. Bars indicate mean ± SD (n = 5)

Biofilm formation by each bacterium was quantified using Crystal Violet staining after culture in YG medium. Absorbance measurements showed that *Paraburkholderia fungorum* exhibited significantly higher absorbance values compared with *Pandoraea norimbergensis* and *C. insecticola* (*P* < 0.05, Mann-Whitney U test with Bonferroni correction), while there was no significant difference between *Pandoraea* and *Caballeronia* (Fig. 5B). These results indicate that *Paraburkholderia* produced a greater amount of biofilm than the other two species under YG culture conditions.

When polymyxin B, an antibiotic that attacks bacterial cell membranes, was added to the YG medium at a concentration of 25 μg/ml, the amount of biofilm produced by *Paraburkholderia fungorum* markedly decreased (Fig. 5B). Under this condition, the absorbance of *Paraburkholderia* was significantly lower than that of *Pandoraea* (P < 0.05, Mann-Whitney U test with Bonferroni correction). The absorbance of *Caballeronia* did not differ significantly from that of either *Pandoraea* or *Paraburkholderia*, showing intermediate values between the two (Fig. 5B). These findings suggest that, in the presence of polymyxin B, *Pandoraea* forms the largest amount of biofilm, whereas *Paraburkholderia* reduces its biofilm production. The concentration of polymyxin B used in this study (25 μg/ml) did not affect bacterial growth (Fig. S4).

## Discussion

Host-microbe symbioses without vertical transmission are characterized by multiple-layered partner choice mechanisms that facilitate the establishment of specific relationships (1). In the well-known legume-rhizobia symbiosis, chemical communication between the host and microbe via nod factors is crucial, and bacterial motility and chemotaxis is important for rapid infection and the formation of specific symbioses (20, 21). In the bean bug *Riptortus pedestris*, once *Caballeronia insecticola* colonize the M4 symbiotic organ, the CR completely closes, preventing secondary infections by other bacteria (14). Additionally, a previous study showed that the infection rate of subsequent bacteria decreases over time, and 18 hours after the oral infection, no subsequent bacteria can colonize the organ (14). This previous study also implies that the time taken to reach the symbiotic organ significantly influences the selection of symbiotic bacteria before establishing the colonization. The results of this study show a significant difference in the time it takes for *C. insecticola* and other bacteria (*Paraburkholderia fungorum* and *Pandoraea norimbergensis*) to reach the symbiotic organ, with *Caballeronia* being the fastest (Fig. 1-3). Moreover, the chemotaxis mutant of *C. insecticola* was slower to reach the symbiotic organ compared to the wild-type symbiont strain (Fig. 1-3), indicating that chemotaxis is crucial for the rapid colonization of the symbiotic organ. Additionally, the co-infection experiments demonstrated that the chemotaxis mutant had reduced competitive ability within the symbiotic organ compared to the wild-type (Fig. 4), strongly suggesting that chemotaxis is a key factor in *Caballeronia* to rapidly and specifically occupy the symbiotic organ. *Caballeronia insecticola* may have acquired chemotactic capabilities through evolutionary process to ensure efficient and swift colonization of the symbiotic organ. To clarify the evolutionary process of the symbiont in more detail, it would be of great interest to investigate what chemical in M4 cues the symbiont-specific chemotaxis.

Microscopic observations revealed that *Pandoraea norimbergensis* formed biofilm-like aggregates within the main duct of the symbiotic organ (Fig. 2 and 5A), which may hinder its colonization of the crypts. *Paraburkholderia fungorum* did not show such aggregates in M4 (Fig. 2 and 5A) but formed clusters within the M3 region (Fig. 1), which seems to cause the delayed colonization of the symbiotic organ. These observations indicate species-specific differences in the patterns and locations of aggregate formation, which may affect colonization efficiency. *In vitro* assays suggested that antimicrobial peptides (AMPs) could influence biofilm formation (Fig. 5B). Within the bug’s gut, the types and levels of AMPs expressed differ between the M3 and M4 regions (5, 22), raising the hypothesis that these spatial variations in host immunity might underlie the distinct aggregate formation patterns observed in *Paraburkholderia* and *Pandoraea*. Although additional experiments will be needed to verify this, our data provide a basis for generating such hypotheses. In contrast, the reduced immune reactivity in *C. insecticola* (23, 24), suggests a divergent strategy in which attenuation of immune resistance, rather than enhancement, facilitates stable symbiosis. Together, these findings imply that both bacterial aggregation and immune interactions could have played key roles in shaping the evolutionary trajectory of the insect–*Caballeronia* symbiosis.

Another notable point is that *C. insecticola* preferentially infects the posterior crypts within M4 (Fig. 3). This infection pattern was absent in the chemotaxis mutant, suggesting the importance of chemotaxis in this process. In some stinkbugs, the posterior crypts of the symbiotic organ play a special role (25). For instance, in the brown-winged green stinkbug, the posterior crypts of the adult’s symbiotic organ become hypertrophied compared to other crypts (26, 27), and the symbionts residing in these crypts are applied to the surface of the eggs (28, 29), ensuring vertical transmission to the next generation. In several stinkbugs associated with *Caballeronia*, vertical transmission of symbionts from eggs and adult feces has been documented (30, 31). Furthermore, after colonization, some of the proliferated bacteria are digested and absorbed by the host in the M4B region (32). Therefore, proliferating in the posterior part of the M4 may allow the bacteria more time before being digested by the host.

Our study revealed that *C. insecticola* uses chemotaxis towards the symbiotic organ to establish an efficient and exclusive symbiotic relationship with its host, *R. pedestris* (Fig. 1-3). This phenomenon was not observed in phylogenetically related bacteria (Fig. 1-3), suggesting that this chemotactic ability was acquired during the evolutionary process of the symbiotic association in the *Caballeronia* lineage. In addition, unlike *Paraburkholderia* and *Pandoraea* species, *Caballeronia* did not form aggregates in M3 and M4 (Fig. 1, 2 and 5), suggesting that the symbiont has some resistance against gut AMPs and/or loss of biofilm-forming capacity in the gut. Although genetic and molecular bases of the chemotaxis and the cell aggregates remain unclear, the dynamic gain and loss of these traits enables *Caballeronia* to specifically associate with the insect host, *R. pedestris*. Our research provides empirical evidence on the ecological trait evolution of symbiotic microorganisms, offering new insights into host-microbe symbioses.

## Acknowledgements

We thank Miyazaki Madoka and Shizuka Ohwaki (AIST) for technical assistance. We are grateful for the helpful comments to Peter Mergaert (CNRS) and Hiroyuki Shimoji (University of the Ryukyus). We also thank Jang Seonghan (KRIBB) for making GFP-labelled bacteria. This work was supported by the Japan Society for the Promotion of Science (JSPS) Research Fellowships for Young Scientists 22KJ0057 and JSPS KAKENHI grants 22H05068.

## Author Contributions

K.I. and Y.K. conceived and designed the study. K.I. conducted experiments and analyzed the data. K.I., A.L. and Y.K. contributed to the final version of the manuscript.

