## Supplemental Figure for "Evolution in spatiotemporal infection patterns of *Burkholderia* sensu lato lineages in the gut of *Riptortus pedestris*"

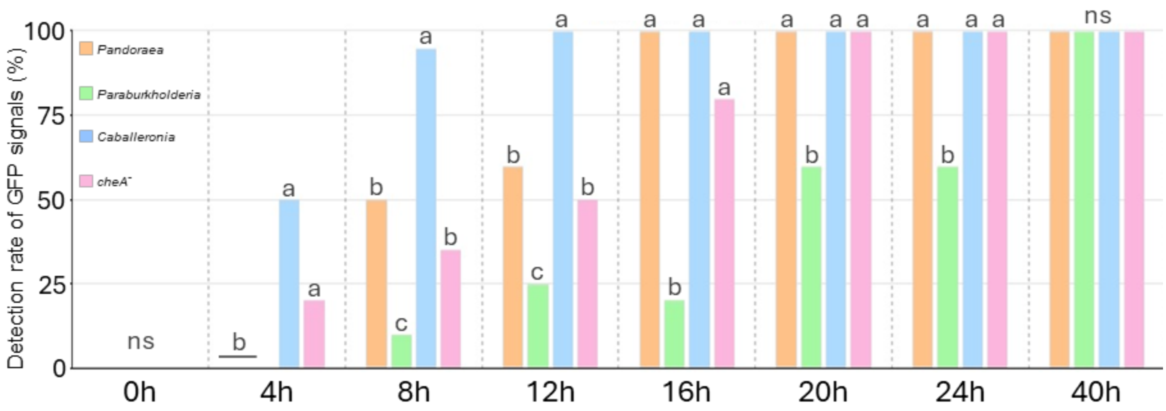


**FIGURE S1. Time to reach the symbiotic organ.**

The Time after inoculation at which GFP signals were detected in symbiotic organs. For each time point and each bacterium, twenty insects were investigated. If even a single bacterial GFP-signal was detected in the main duct of the M4 region, it was counted as “positive”. The different letters indicate statistically significant differences ($P<0.05$). The statistical significances were analyzed by The Fisher’s exact test with Bonferroni correction.

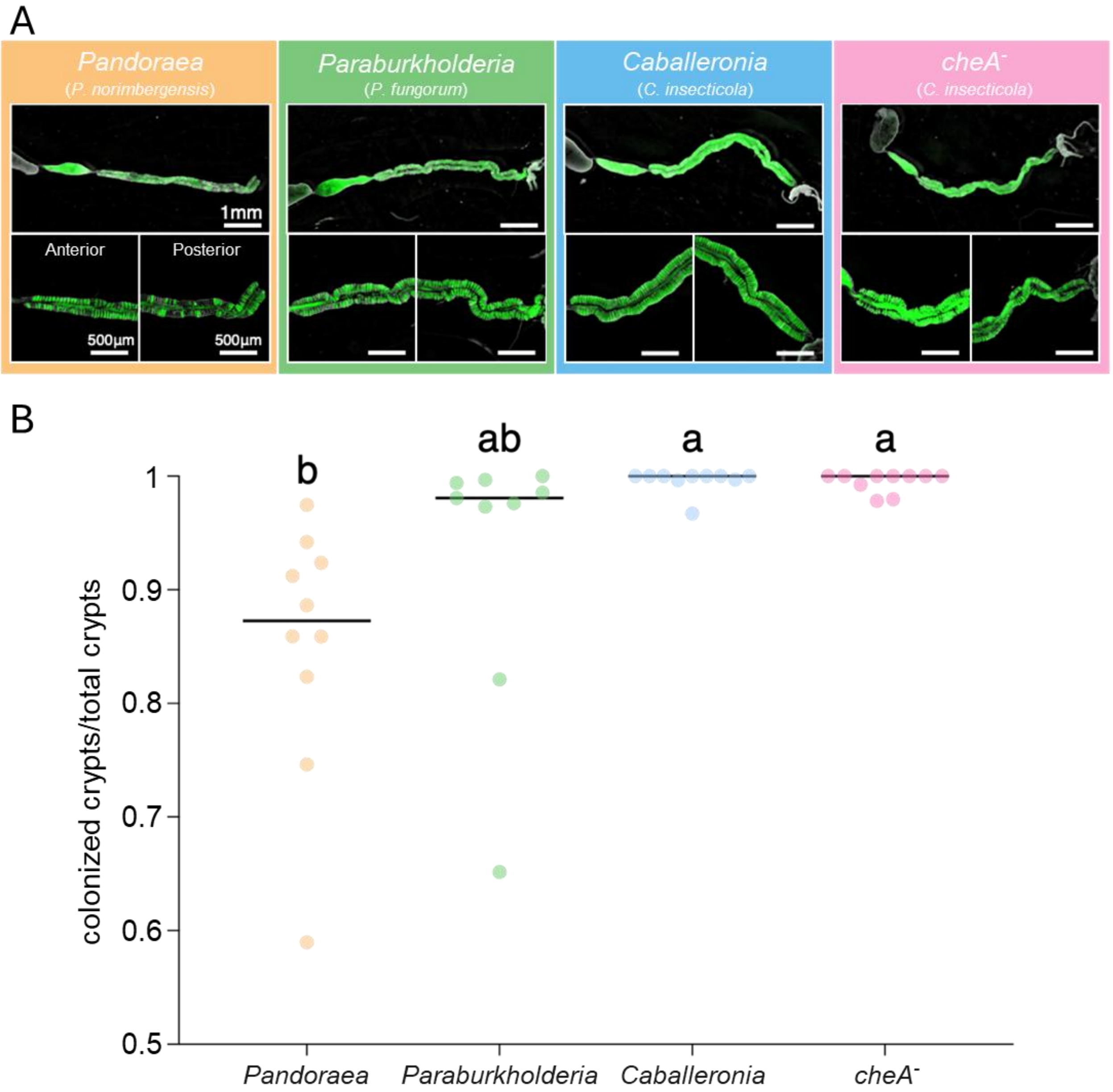


**FIGURE S2. Colonization frequency of crypts 5 days after oral inoculation.**
(A) Images of the GFP-labeled bacteria in symbiotic organ. (B) The rate of colonized crypts by bacteria. The different letters indicate statistically significant differences ($P<0.05$) analyzed by The Mann-Whitney U test with Bonferroni correction.

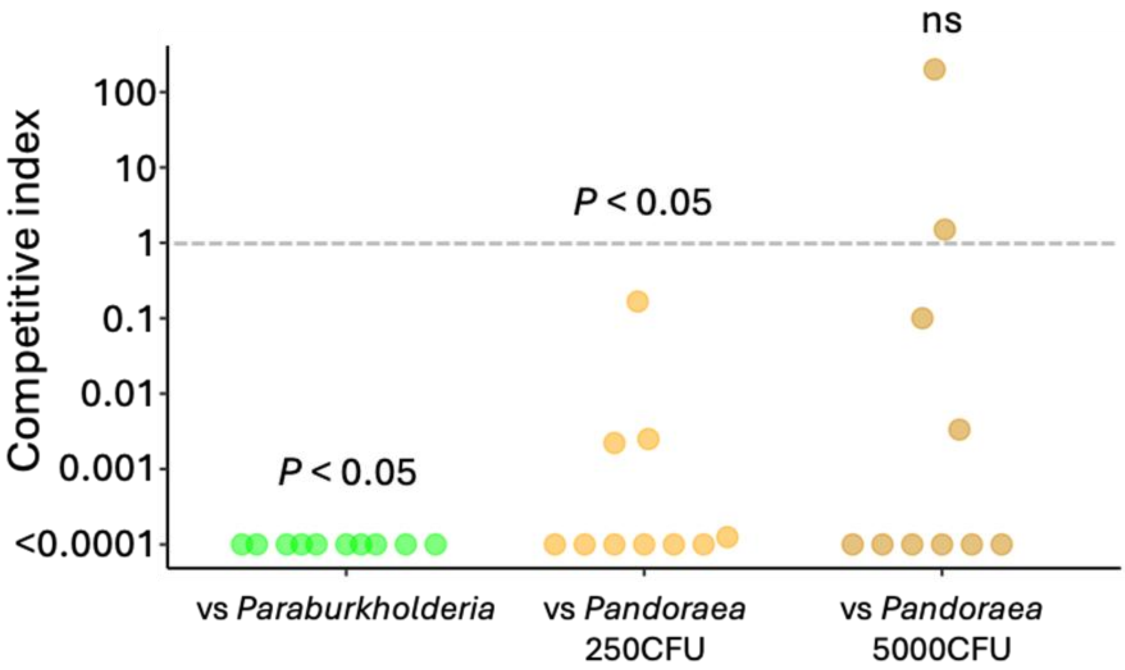


**FIGURE S3. Competitiveness of *Caballeronia insecticola*.**

Competitiveness of *C. insecticola* within symbiotic organs. Competitive index values were obtained by (output competitor CFU/input competitor CFU)/(output *C. insecticola* CFU/input *C. insecticola* CFU) and statistically evaluated by the 1-sample t test (against CI = 1.0).

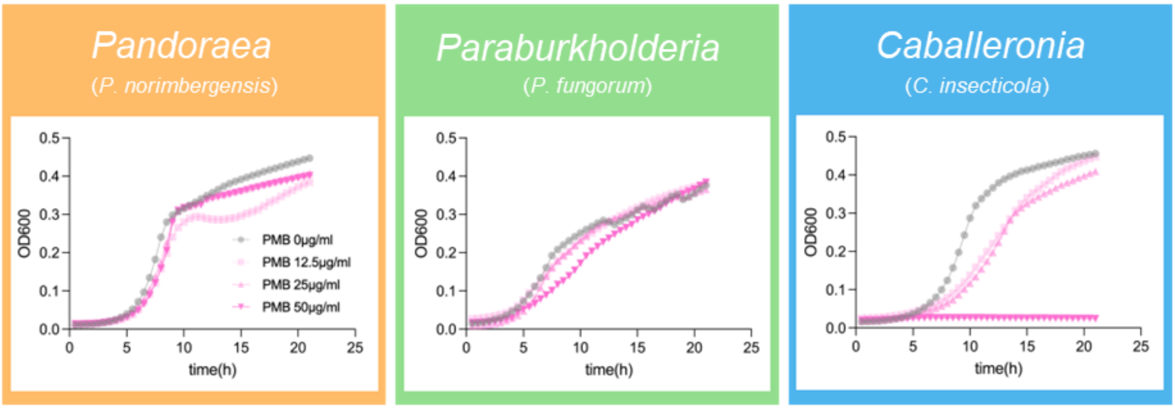


**FIGURE S4. Growth inhibition of Polymyxin B.**

Growth curves of the *Pandoraea norimbergensis*, *Paraburkholderia fungorum*, *Caballeronia insecticola* with different concentrations of polymyxin B. For each bacterial strain and each polymyxin B concentration (0, 12.5, 25, and 50 μg/mL), four independent replicates were measured.
